# Position-specific secondary acylation determines detection of lipid A by murine TLR4 and caspase-11

**DOI:** 10.1101/2021.12.16.472973

**Authors:** Erin M. Harberts, Daniel Grubaugh, Daniel C. Akuma, Sunny Shin, Robert K. Ernst, Igor E. Brodsky

## Abstract

Immune sensing of the Gram-negative bacterial membrane glycolipid lipopolysaccharide (LPS) is both a critical component of host defense against Gram-negative bacterial infection, and a contributor to hyper-inflammatory response, leading to sepsis and death. Innate immune activation by LPS is due to the lipid A moiety, an acylated di-glucosamine molecule that can activate inflammatory responses via the extracellular sensor TLR4/MD2 or the cytosolic sensor caspase-11 (Casp11). The number and length of acyl chains present on bacterial lipid A structures vary across bacterial species and strains, which affects the magnitude of TLR4 and Casp11 activation. TLR4 and Casp11 are thought to respond similarly to various lipid A structures, as tetra-acylated lipid A structures do not activate either sensor, whereas hexa-acylated structures activate both sensors. However, direct analysis of extracellular and cytosolic responses to the same sources and preparations of LPS/lipid A structures have been limited, and the precise features of lipid A that determine the differential activation of each receptor remain poorly defined. To address this question, we used rationally engineered lipid A isolated from a series of bacterial acyl-transferase mutants that produce novel, structurally defined molecules. Intriguingly, we find that the location of specific secondary acyl chains on lipid A resulted in differential recognition by TLR4- or Casp11, providing new insight into the structural features of lipid A required to activate either TLR4- or Casp11. Our findings indicate that TLR4 and Casp11 sense non-overlapping areas of lipid A chemical space, thereby constraining the ability of Gram-negative pathogens to evade innate immunity.

## Introduction

Innate immune receptors alert the host to infection through sensing of pathogen-associated molecular patterns (PAMPs). Lipopolysaccharide (LPS) and lipooligosaccharide (LOS) are essential structural components of Gram-negative bacterial membranes that activate the extracellular membrane-bound Toll-like receptor 4 (TLR4)/MD-2 signaling complex (hereafter called TLR4) and the intracellular receptors human caspase-4/5 and mouse caspase-11 [1]. Mice lacking either TLR4 or Casp11 are highly susceptible to Gram-negative bacterial infection and conversely, are resistant to LPS-driven models of acute sepsis [2-5]. LPS is composed of O-antigen, a highly variable glycosyl structure attached to a more conserved core oligosaccharide, which together are anchored into the outer leaflet of the outer membrane by the highly conserved lipid A moiety, whereas LOS contains the core oligosaccharide portion and lacks O-antigen. Lipid A, the membrane anchor of LPS/LOS, is sensed by both TLR4 and caspase-11/4/5, and is thus the primary driver of endotoxic shock during Gram-negative bacterial infection [6, 7]. Lipid A is composed of a β(1→6) linked di-glucosamine backbone, two 3-OH acyl groups attached at each of the N-linked (2 and 2’) and O-linked (3 and 3’ position) positions of the di-glucosamine backbone, and two terminal phosphate moieties at the 1 and 4’ positions. The acyl chains bound directly to the glucosamine sugars are the primary acyl chains and can in turn be modified with addition of secondary acyl chains via ester linkages [8]. The number, position, and length of the acyl chains vary widely across different species and strains of bacteria, as well as in response to environmental conditions [9]. Importantly, these structural differences have a direct effect on the magnitude and duration of the innate immune response, but the precise effect of specific acyl chain additions/removals has not yet been described [10].

Extracellular LPS is detected by the plasma membrane-localized TLR4/MD2 receptor signaling complex. When not bound by a ligand, TLR4 exists as a monomer spanning the plasma membrane with extra- and intracellular domains. Receptor complex initiation begins when LPS is bound by lipopolysaccharide-binding protein (LBP), which then transfers LPS to CD14 and subsequently the MD-2 co-receptor. MD-2 presents LPS to TLR4, causing a conformational change that enables dimerization of TLR4 monomers. This brings the TLR4 cytoplasmic signaling domains into proximity with each other and propagates downstream signaling [11, 12]. The structure of the lipid A determines binding to the TLR4/MD-2 complex [13, 14]. In general, hexa-acylated lipid A structures are known agonists of TLR4, whereas tetra-acylated lipid A is antagonistic [15]. In addition to acyl chain numbers, lipid A acyl chain length has an impact on downstream signal strength. This has been shown previously in mutants of *Haemophilus influenzae, Neisseria meningitidis, Neisseria gonorrhoeae*, and *Salmonella typhimurium* lacking the secondary lipid A acyltransferase HtrB/LpxL, resulting in a penta-acylated lipid A lacking C14:0 (myristate), leading to reduced inflammatory potential and virulence [16-22]. In addition, the acylation state of lipid A, alterations in the terminal phosphate moieties can alter innate immune recognition. For example, the lipid A phosphatases, LpxE and LpxF from Francisella species, are site-specific enzymes that remove either the 1- or 4’-postion phosphate groups, respectively, and affect recognition by TLR4 and resistance to cationic antimicrobial molecules [23-29]. Notably, while both human and mouse TLR4 detect extracellular LPS, they exhibit structural-dependent differences in their responsiveness to distinct lipid A structures, as the human MD2/TLR4 complex is less responsive than mouse TLR4 to under-acylated forms of lipid A [30, 31]. Furthermore, differential TLR4 activation between temperature-regulated lipid A structures, synthesized by *Y. pestis* is more pronounced in human cells, which do not recognize tetra-acylated lipid A, as compared to mouse cells, which maintain limited recognition of tetra-acylated lipid A [32] and highlights the selectivity of the structure-activity relationship (SAR) of lipid A with the TRL4/MD-2 complex.

While TLR4 senses the presence of extracellular LPS, the intracellular proteins caspase-11 (Casp11) in mice, and its human orthologs CASP4 and CASP5 detect LPS that enters the cytosol due to vacuolar escape by cytosolic pathogens such as *Burkholderia pseudomallei* [33] or disruption of the vacuolar membrane by bacterial or cellular factors [4, 7, 34, 35]. Unlike TLR4, caspases do not strictly require a coreceptor for detecting LPS, although recent studies have revealed an important accessory role for guanylate binding proteins (GBPs) in full activation of this pathway [36-39]. Briefly, lipid A binds directly to the caspase activation and recruitment domain (CARD) of Casp11, leading to its oligomerization [7], followed by proximity-induced autoprocessing and formation of the non-canonical inflammasome [40]. Activated Casp11 subsequently cleaves downstream effector molecules, including Gasdermin D (GSDMD), leading to formation of GSDMD pores in the plasma membrane that mediate pyroptotic cell death and release of interleukin-1 (IL-1) family cytokines and intracellular alarmins [41, 42].

While the structural requirements of lipid A bound to MD-2 and TLR4 are better understood [43], the structural requirements and the contribution of individual lipid A components to Casp11 activation are undefined. Furthermore, because lipid A structure varies greatly among bacterial species and very little direct comparison of TLR4 and Casp11 responsiveness to individual lipid A structures exist, we have only a limited understanding of how distinct lipid A structures activate these two endotoxin sensing pathways.

Here, we used defined mutants of *Yersinia pestis* that lack one or both of the secondary acyltransferases, MsbB/LpxM and LpxP to generate defined structures of bacterial lipid A, allowing us to directly dissect the contribution of specific lipid A secondary acyl-chain modifications to activation of murine TLR4 and Casp11. *Y. pestis* only synthesizes LOS due to a mutation in O-antigen synthesis. Intriguingly, we found that in murine cells, position-specific addition of the fifth acyl chain to lipid IVa by either MsbB or LpxP acyltransferases converts the non-activating tetra-acylated lipid IVa to a specific agonist of either Casp11 or TLR4, respectively. Collectively, these findings indicate that mouse Casp11 and TLR4 are differentially activated by structurally distinct forms of lipid A. In particular, the location of secondary acyl chains differentially impacts the ability of these two lipid A sensors to detect various lipid A species. These findings highlight the overlapping constraints that these innate immune LPS sensing molecules place on the lipid A structures of pathogens.

## Methods

### Bacterial strains and lipid A extraction

Bacterial strains used in these studies were created in the laboratory of Dr. B. Joseph Hinnebusch and are previously described in Rebeil *et al*., 2006 [44]. Bacteria were cultured in BHI media (BD #237200) prepared according to manufacturer’s recommendation and supplemented with 1 mM MgCl_2_. Briefly, overnight cultures (10ml) were grown shaking (200 rpm) at 26L and used to inoculate 1L liquid cultures which were grown for 18-24 hours under the same conditions [45]. Bacterial pellets were harvested by centrifuging at 10,000 x g for 20 minutes.

Extraction of LOS was performed as previously described [45] with modifications noted below. A hot phenol extraction was then performed by resuspending the resultant pellet from 1L of culture in 60mL endotoxin free water. This suspension was split evenly among three 50mL conical tubes and an equal volume (20mL) of 90% phenol was added to each tube which were incubated at 65□ for 1 hour with intermittent vortexing. The mixture was cooled on ice for 5 minutes and centrifuged at 3,000 x g for 20 minutes at room temperature (RT). The aqueous (top) layer was removed and placed into a new 50mL conical. An equal volume of endotoxin-free water was added to the original tube and the extraction steps repeated. The two aqueous layers were combined and dialyzed using 1kD MWCO tubing (Spectrum Laboratories #132105) for 24-36 hours at 4□ in at least 4L of ddH_2_O. The ddH_2_O was changed several times during dialysis period. To remove any remaining insoluble material, the aqueous solution was centrifuged at 5,000 x g for 20 minutes at room temperature (RT) and the solution lyophilized. The dried material was resuspended in 10mL of 10mM Tris-HCL (pH 7.4), then 25ug/mL RNase (Qiagen #19101) and 100ug/mL of DNase I (Roche #10104159001) were added followed by a 2 hour incubation at 37□. 100ug/mL of Proteinase K (Invitrogen #25530-015) were then added and the incubation at 37□ continued for an additional 2 hours. 5mL of water saturated phenol were added before the solution was mixed via vortex and then centrifuged at 3,000 x g for 20 minutes at RT. The resulting aqueous fraction was dialyzed for at least 12 hours, insoluble fraction removed, and lyophilized as before, resulting in extracted bacterial LOS. The product was further purified as previously described using chloroform/methanol washes to ensure removal of hydrophilic lipids and re-extraction to remove any remaining lipoproteins [45, 46]. Mild acid hydrolysis was carried out as previously described to further convert LOS to lipid A [45].

### Cell culture

Reporter cell lines hTLR4 HEK-Blue (Invivogen) and RAW-Blue (Invivogen) were cultured in DMEM supplemented with 10% fetal bovine serum, 100U/mL Penicillin-Streptomycin, and 2mM L-Glutamine in a humidified tissue culture incubator at 37□ and 5% CO_2_. Cells were split into a 96-well flat bottom plate, allowed 24-48 hours to adhere, and then stimulated with agonists over a 5-log concentration range. After 18 hours of culture with agonist, 20μL of cell culture supernatant were added to 180 μL of Quanti-Blue detection media (Invivogen) and developed in the tissue culture incubator for 20 minutes. Optical density (OD) at 620 nm was determined using a DTX 880 multimode plate reader (Beckman Coulter, Brea, CA). Mouse bone marrow-derived macrophages were cultured in DMEM supplemented with 10% fetal bovine serum, 10mM HEPES, 1mM sodium pyruvate, and 30% L929 cell-conditioned media. Cells were grown for 6-7 days in petri plates before reseeding into 48-well plates at 16-20 hours before LOS or lipid A treatment.

### LOS and lipid A intracellular delivery

For LOS transfection, mouse bone marrow-derived macrophages were primed 4 hours with 400ng/mL Pam3CSK4 in DMEM + 10% L929 supernatant and then transfected with the indicated doses of endotoxin with 0.2% FuGENE HD in Opti-MEM. Twenty hours after transfection, cell supernatants were assayed for cell death and cytokine concentrations. *L. monocytogenes-*mediated delivery of LOS was performed as previously described [34]. Briefly, BMDMs were primed with 1µg/mL HMW poly(I:C) for 16 hours, then infected with *L. monocytogenes* strain 10403S at a MOI of 5 in the presence of LOS. One hour after infection, media was replaced with media containing 20µg/mL gentamicin. Four hours after infection, cell supernatants were harvested to quantify cell death. For both methods of intracellular delivery, cell death was assayed using a lactate dehydrogenase release assay kit (Takara BioProduct) according to manufacturer’s instructions.

### IL-1β ELISA

Supernatants were applied, along with recombinant cytokine standards, to Immulon ELISA plates (ImmunoChemistry Technologies) that were pre-coated with anti-IL-1β capture antibody (eBioscience). IL-1β was then detected with a secondary biotinylated antibody (eBioscience) followed by streptavidin conjugated to horseradish peroxidase (BD Biosciences). Peroxidase activity was detected by o-phenylenediamine hydrochloride (Sigma-Aldrich) in citrate buffer. Reactions were stopped with sulfuric acid and read at 490nm.

### Graphs and statistics

Data were graphed and statistical analyses performed in GraphPad Prism 8.4.3 (San Diego, California). Statistically significant differences between experimental conditions were determined by multiple t tests using the Holm-Šídák method. Best fit lines were calculated in Prism using an equation for 4-parameter variable slope regressions.

## Results

### Distinct penta-acylated lipid A species enable defined analysis of extracellular and cytosolic innate immune responses to endotoxin

Lipid A structure varies broadly among bacterial species [9], many of which undergo regulated changes to their lipid A in response to environmental conditions, including temperature and pH [47]. In particular, *Yersinia pestis* (*Yp*) grown at an environmental temperature (26□) produces a hexa-acylated lipid A, whereas lipid A prepared from *Yp* grown at mammalian body temperature (37□) is tetra-acylated due to the absence of secondary acyl chains [32, 48]. *Yp* lipid A structure determines how the bacteria are recognized by the host and correlates with pathogenicity [49]. When grown at mammalian temperature, *Yp* evades immune recognition by TLR4, an adaptation that allows for evasion of innate immune defense and enhances virulence in mammals [50, 51]. In *Yp*, the addition of the secondary C12 and C16:1 acyl chains is mediated by the acyltransferases MsbB and LpxP, respectively (**Figure 1**) [44]. Thus, loss of LpxP results in absence of the O-linked 16:1 acyl-chain on the 2’ 14-carbon acyl-chain, whereas deletion of *msbB* results in the loss of the O-linked 12-carbon acyl-chain on the 3’ 14-carbon acyl-chain.

**Figure 1.**
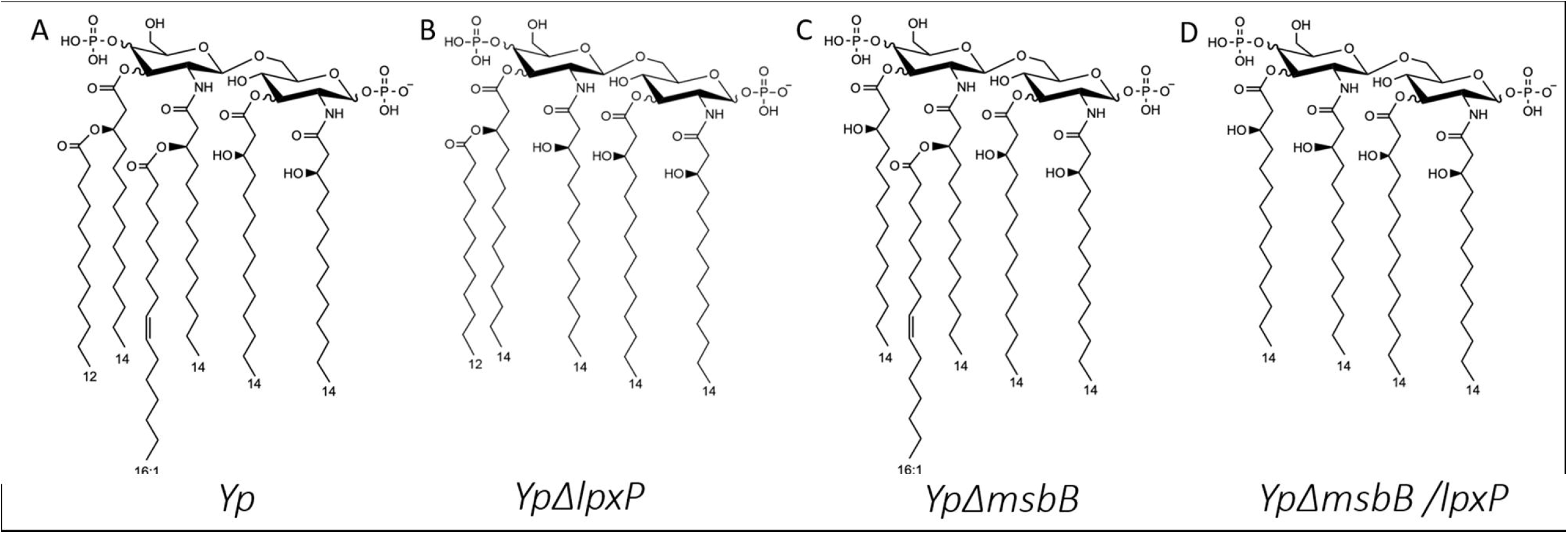
Chemical structures of lipid A molecules. Structures are shown for WT *Yersinia pestis* (A), *Yp*Δ*lpxP* (B), *Yp*Δ*msbB* (C), and *Yp*Δ*lpxP/msbB* (D) lipid A, all grown at 26L. When describing these molecules, the 1-position is where the phosphate group on the right side of the molecule is added with the carbons in the ring sequentially numbered moving down and to the left. Similarly, the 1’-position is where the link is made between the two carbon rings again numbered sequentially with the 2’-poisiton being the next carbon down and to the left. The second phosphate group is added at the 4’-position.

Deletion of both *msbB* and *lpxP* results in a 14-carbon tetra-acylated lipid A. While hexa-acylated lipid A structures are detected by both TLR4 and Casp11, and tetra-acylated lipid A structures are not detected by either, penta-acylated LPS structures are variable with respect to their ability to activate TLR4 and Casp11 [4, 8, 52, 53]. The underlying basis for selective responses of intra- and extracellular LPS sensing pathways to different LPS structures is therefore not clear. Moreover, responses of murine and human extra- and intra-cellular LPS sensing pathways have been studied in response to similar lipid A structures isolated from different bacterial species under different growth conditions, thus limiting the ability to make direct comparisons between murine and human systems. We previously established a system in

*Y. pestis* to purify defined LOS structures that have distinct abilities to stimulate TLR4-dependent responses [45]. Here, we took advantage of this approach to produce LOS structures that are identical other than the positioning and number of secondary acyl chains (**Figure 1**), thereby enabling us to define the innate immune responses to closely related penta-acylated lipid A structures in murine and human system (Alexander-Floyd, Harberts, et al., companion manuscript).

### LpxP-mediated lipid A modifications allow for TLR4 activation

To dissect the contribution of specific acyl chain lipid A modifications to TLR4 signaling, the HEK-Blue mTLR4 reporter cell line was used to define TLR4-dependent signaling in response to LOS and lipid A isolated from WT *Yersinia pestis* (*Yp*) and acyltransferase mutant strains lacking one or both of *lpxP* and *msbB*. Notably, LOS isolated from the *Yp* Δ*msbB* mutant, which lacks the secondary C12 acyl chain at the 3’ position, activated TLR4 signaling in HEK-Blue mTLR4 cells similarly to the pro-inflammatory WT *Yp* hexa-acylated structure, whereas penta-acylated lipid A derived from *Yp* Δ*lpxP*, lacking the secondary C16:1 acyl chain at the 2’ position was a poor TLR4 activator, similar to tetra-acylated lipid A isolated from the *Yp* Δ*msbB/lpxP* double mutant (**Figure 2A**). This was the case for both lipid A and corresponding LOS molecules, though WT *Yp* LOS showed higher activity, indicating that the oligosaccharide moieties beyond lipid A do not fundamentally affect TLR4 recognition.

**Figure 2.**
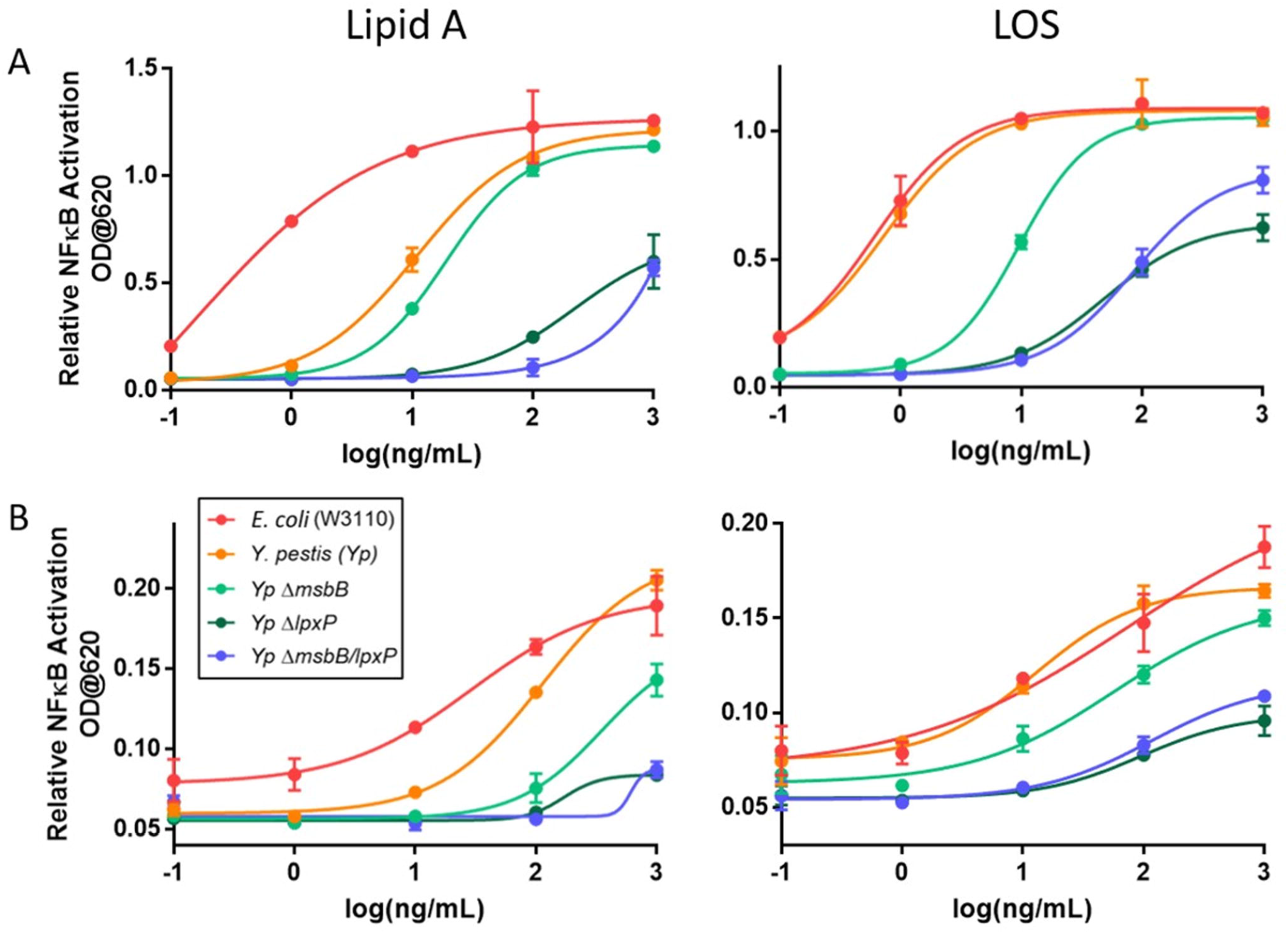
LPS and lipid A from *Yersinia pestis* Δ*msbB* but not Δ*lpxP* agonize TLR4. LPS/LOS and lipid A extracted from *E. coli* (red), WT *Yp* (orange), *Yp* Δ*msbB* (light green), *Yp* Δ*lpxP* (dark green), and *Yp* Δ*msbB*/*lpxP* (blue) are cultured with HEK-Blue mTLR4 cells (A) or RAW-Blue cells (B) for 18 hours. SEAP reporter is then detected in cell culture supernatant and OD at 620 nm, indicative of relative NFκB activation, is graphed over the 5-log concentration range tested. Average ± SD of biological duplicates for each data point is shown, graph is representative data from three independent experiments producing consistent results.

Consistently, RAW-Blue cells, which express an NF-κB-driven secreted alkaline phosphatase, also exhibited elevated activation in response to LOS and lipid A from *Yp* Δ*msbB* compared to WT Yp-or *Yp* Δ*lpxP*-derived lipid A and LOS (**Figure 2B**), though the differences in activation by different TLR4 ligands were not as pronounced in the RAW-Blue cell line. This may be due to differences in the level of endogenous TLR4 expression in RAW macrophage-like cells in comparison to HEK293 cells. These findings are consistent with previous observations that penta-acylated LPS structures containing acyl chains of 16 carbons or longer are generally weak agonists for TLR4/MD-2 due to their poor fit within the MD-2 binding pocket. Together, these findings identify a previously unknown minimum stimulatory component of lipid A signaling through TLR4 and demonstrate that penta-acylated lipid A containing a secondary acyl-chain at the 2’ position, but not the 3’ position is sufficient to activate TLR4 signaling.

### MsbB-dependent acylation of lipid A is required for activation of Casp11-dependent cell death

Caspase-11 (Casp11) is the intracellular sensor of lipid A in mice. While both TLR4 and Casp11 sense lipid A, the structural requirements for Casp11-mediated detection of lipid A are poorly defined. To understand the role of specific lipid A modifications in Casp11 activation, we primed wild-type (C57BL/6) and Casp11-deficient (*Casp11*^*-/-*^) mouse bone marrow-derived macrophages (BMDMs) with Pam3CSK4 for four hours followed by transfection with LOS containing the defined lipid A structures in **Figure 1**. We assessed the ability of distinct lipid A structures to activate Casp11 by measuring Casp11-dependent cell death using a lactate dehydrogenase release assay (LDH). As expected, hexa-acylated lipid A induced cell death in WT BMDMs relative to *Casp11*^*-/-*^ BMDMs at each of the concentrations tested (**Figure 3A**), whereas tetra-acylated lipid A from *Yp* Δ*msbB*/*lpxP* did not (**Figure 3B**). However, penta-acylated lipid A derived from *Yp* Δ*msbB*, which lacks a C12 secondary acyl chain on the 3’ O-linked primary acyl group (**Figure 1**), induced minimal levels of cell death (**Figure 3C**). This finding is consistent with prior observations that penta-acylated lipid A structures, particularly those containing C16-length acyl chains, are generally weak activators of Casp11 or are Casp11 antagonists [4], and contrasts with the ability of lipid A derived from *Yp* Δ*msbB* to activate TLR4.

**Figure 3:**
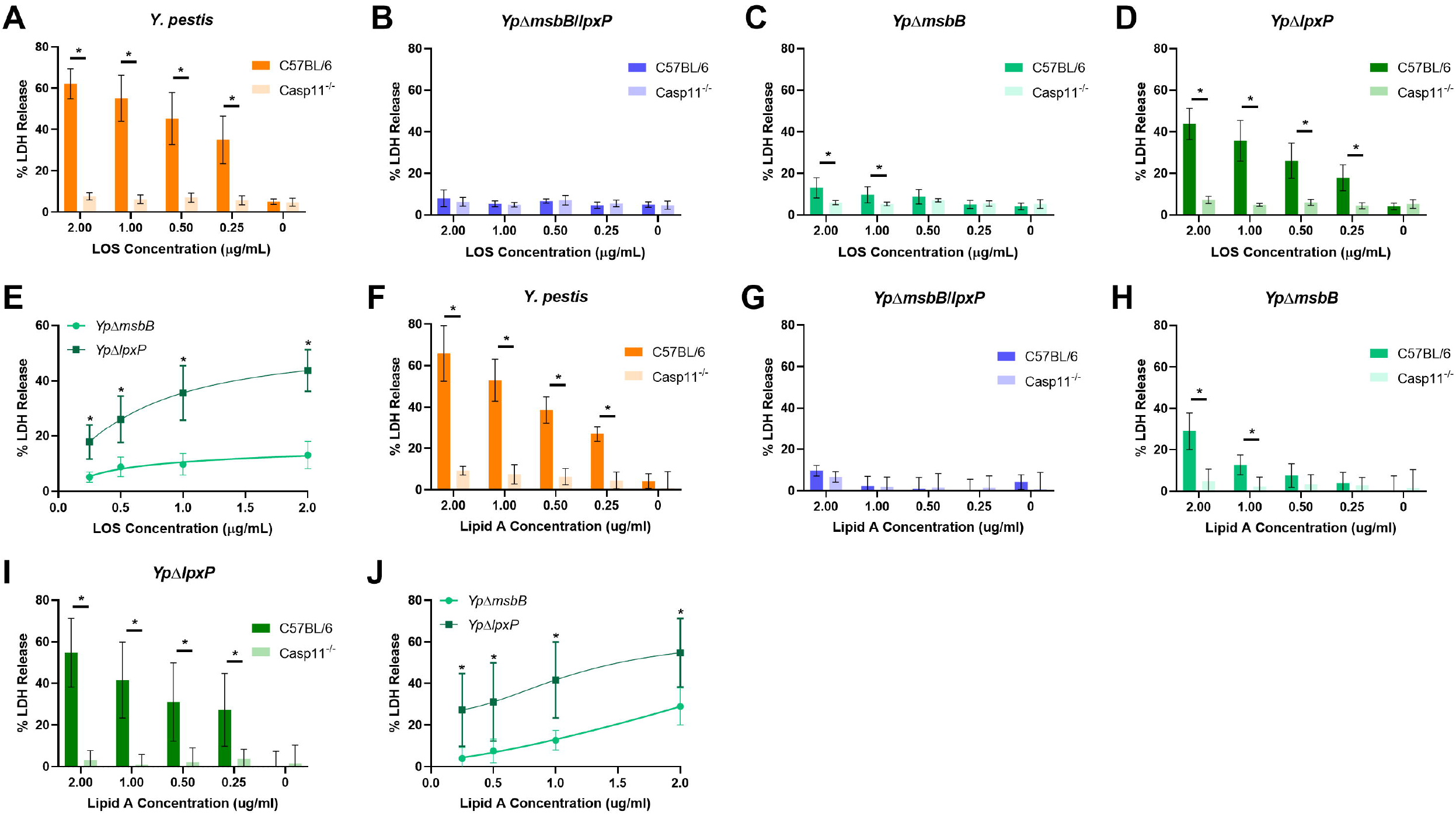
Secondary acyl chain position determines the ability of penta-acylated LPS to induce Casp11-dependent cell death. Pam3CSK4 primed BMDMs from wild-type (C57BL/6) and Casp11-deficient (Casp11^-/-^) mice were transfected with the indicated concentrations of LOS (A,C) or lipid A (B,D). Twenty hours after transfection, cell supernatants were assayed for lactate dehydrogenase activity. Lactate dehydrogenase activity was determined relative to cells lysed using Triton X-100. Data are presented averages (+/- SD) of three independent experiments performed in triplicate. A-B: *, significant difference between C57BL/6 and Casp11^-/-^, p<0.05, t-test using Holm-Sidak method for multiple hypothesis testing. C-D: *, significant difference between *Yp*Δ*msbB* and *Yp*Δ*lpxP*, p<0.05, t-test using Holm-Sidak method for multiple hypothesis testing.

Intriguingly, penta-acylated lipid A derived from *Yp* Δ*lpxP* and therefore lacking the secondary acyl group linked to the 2’ amino group acyl chain (**Figure 1**), induced significantly increased levels of cell death in WT BMDMs compared to *Casp11*^*-/-*^ BMDMs, and nearly the same extent of cell death as hexa-acylated LPS (**Figure 3D**). Additionally, *Yp* Δ*lpxP* LOS induced significantly higher levels of cell death than *Yp* Δ*msbB* across a wide dose range (**Figure 3E**), indicating that a single acyl chain position alters the capacity of penta-acylated LOS to activate Casp11. Notably, the same pattern of differential stimulatory capacity was observed when co-administered with *Listeria monocytogenes* [34] (**Supplemental Figure 1**), indicating that differences in transfection efficiency are unlikely to account for the difference observed in their stimulatory capacity. Altogether, these findings suggest that the presence or absence of secondary acyl chain modifications is a major determinant of Casp11 activation.

Previous studies using LPS from *Helicobacter pylori* and *Rhizobium galegae*, which also produce primarily a penta-acylated lipid A structure, found that these molecules inhibit activation of Casp11 by agonist LPS when co-transfected into BMDMs. In contrast, lipid IVa, a structure similar to *Yp* Δ*msbB*/*lpxP* lipid A, does not block Casp11 activation by agonist LPS [4]. To test whether *Yp* Δ*msbB* lipid A could also block Casp11 signaling, we transfected BMDMs with increasing concentrations of *Yp* Δ*msbB* or *Yp* Δ*msbB*/*lpxP* LOS or *R. sphaeroides* (LPS-*Rs*) LPS in the presence of *Salmonella* Minnesota LPS, a potent Casp11 agonist [34]. As expected, tetra-acylated *Yp* Δ*msbB*/*lpxP* LOS failed to inhibit cell death in response to Casp11 agonist LPS, whereas LPS-*Rs*, which binds but does not activate Casp11 strongly blocked cell death at low concentrations. Despite being penta-acylated, we found that *Yp* Δ*msbB* LOS acted more like the *Yp* Δ*msbB*/*lpxP* LOS, partially blocking Casp11 activation at only at the highest concentrations (**Supplemental Figure 2**). These data indicate that acyl chain number alone does not determine Casp11 antagonism.

LOS contains an oligosaccharide moiety which could impact Casp11-dependent innate responses. To test whether the difference in Casp11 activation levels between the various LOS molecules was directly attributable to differences in lipid A acylation, we assayed Casp11-dependent cell death in response to cytosolic delivery of purified lipid A lacking core oligosaccharide from each strain. Consistently, both *Yp* and *Yp* Δ*lpxP* lipid A structures induced robust cell death, whereas lipid A from *Yp* Δ*msbB* largely failed to induce detectable cell death except at the highest concentrations, and tetra-acylated lipid A derived from *Yp* Δ*msbB*/*lpxP* failed to induce cell death entirely (**Figure 3E-H**). As with their LOS structural counterparts, *Yp* Δ*lpxP*-derived lipid A again induced significantly more cell death that lipid A derived from *Yp* Δ*msbB* (**Figure 3J**), indicating that the number and position of acyl chains is the primary determinant for Casp11 activation by bacterial LOS or lipid A.

### MsbB-dependent acylation of lipid A is required for activation of Casp11-dependent IL-1β secretion

Activation of the Casp11 non-canonical inflammasome also triggers the processing and release of IL-1 family cytokines via secondary activation of NLRP3 [35]. To directly test the contribution of acyl chain position to IL-1β release, we assessed IL-1β levels in the supernatants of BMDMs transfected with LOS variants. As expected, LOS from WT *Yp* induced significant IL-1β release at all four tested concentrations (**Figure 4A)**, while tetra-acylated LOS purified from *Yp* Δ*msbB*/*lpxP* bacteria failed to induce significant IL-1β release relative to *Casp11*^*-/-*^ control BMDMs (**Figure 4B**). LOS derived from the *Yp* Δ*msbB* (**Figure 4C**) mutant led to minimal IL-1β release, as compared to *Yp* Δ*lpxP* **(Figure 4D)**, which induced significant Casp11-dependent IL-1β secretion (**Figure 4E**). Similar results were observed following transfection of purified lipid A, with WT *Yp* and *Yp* Δ*lpxP* lipid A leading to IL-1β release at all concentrations and *Yp* Δ*msbB* lipid A inducing IL-1β secretion when transfected at high concentrations (**Figure 4F-J**). Taken together, these data indicate that Casp11 inflammasome response to penta-acylated LOS/lipid A requires a secondary O-linked acylation at the 3’ acyl chain, and that, similar to TLR4, the position of secondary acylation plays a critical role in determining the ability to activate Casp11.

**Figure 4:**
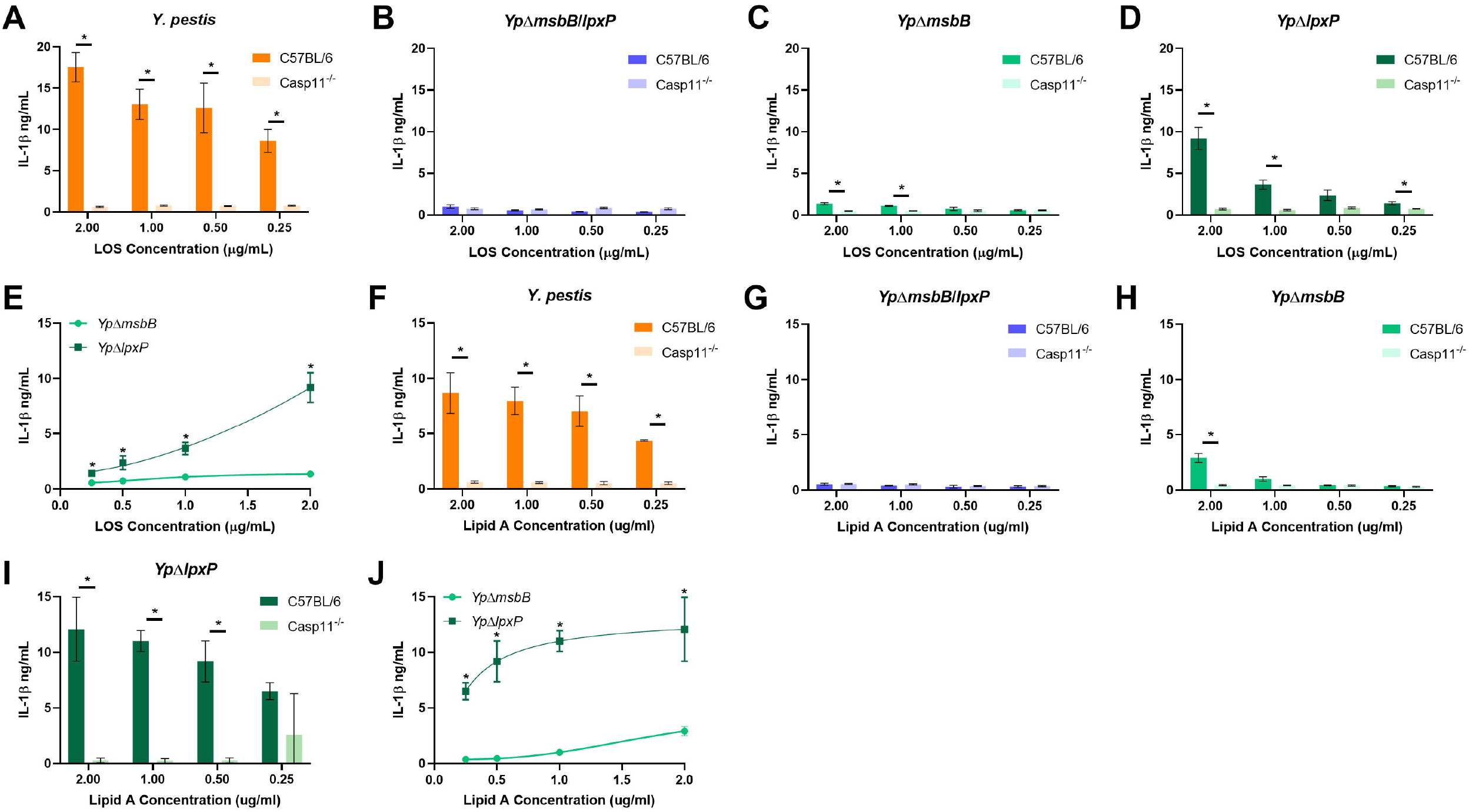
Secondary acyl chain positions contribute to the ability of penta-acylated LPS to induce IL-1β secretion. Pam3CSK4 primed BMDMs from wild-type (C57BL/6) and Casp11-deficient (Casp11^-/-^) mice were transfected with the indicated concentrations of LOS (A,C) or lipid A (B,D). Twenty hours after transfection, cell supernatants were collected and IL-1β concentration was measured using ELISA. Data are representative of three independent experiments performed in triplicate. A-B: *, significant difference between C57BL/6 and Casp11^-/-^, p<0.05, t-test using Holm-Sidak method for multiple hypothesis testing. C-D: *, significant difference between *Yp*Δ*msbB* and *Yp*Δ*lpxP*, p<0.05, t-test using Holm-Sidak method correction for multiple hypothesis testing.

### Guanylate binding proteins on Chromosome 3 are dispensable for differential sensing of penta-acylated LOS

Guanylate Binding Proteins (GBPs) play an important role in facilitating Casp11/Casp4-dependent responses to cytosolic LPS in murine and human macrophages [36, 38, 54-57]. In particular, human GBP1 was recently shown to directly bind and detect LPS [37]. Since human GBP1 shares sequence homology with several mouse GBPs present on chromosome 3 [58], we considered the possibility that distinct LOS or lipid A species may exhibit differential requirement for GBPs in activation of Casp11. To test this possibility, we transfected BMDMs derived from B6 and *Gbp*^*Chr3-/-*^ mice, which lack all of the GBPs on mouse chromosome 3 [59] with the hexa-, tetra-, and penta-acylated LOS structures as described above. *Gbp*^*Chr3-/-*^ BMDMs did not display a significant loss of inflammasome activity in response to either WT or *Yp* Δ*lpxP* Yp LOS (**Figure 5**), indicating that the GBPs on chromosome 3 do not differentially control responsiveness to different penta-acylated LOS structures under these conditions.

**Figure 5:**
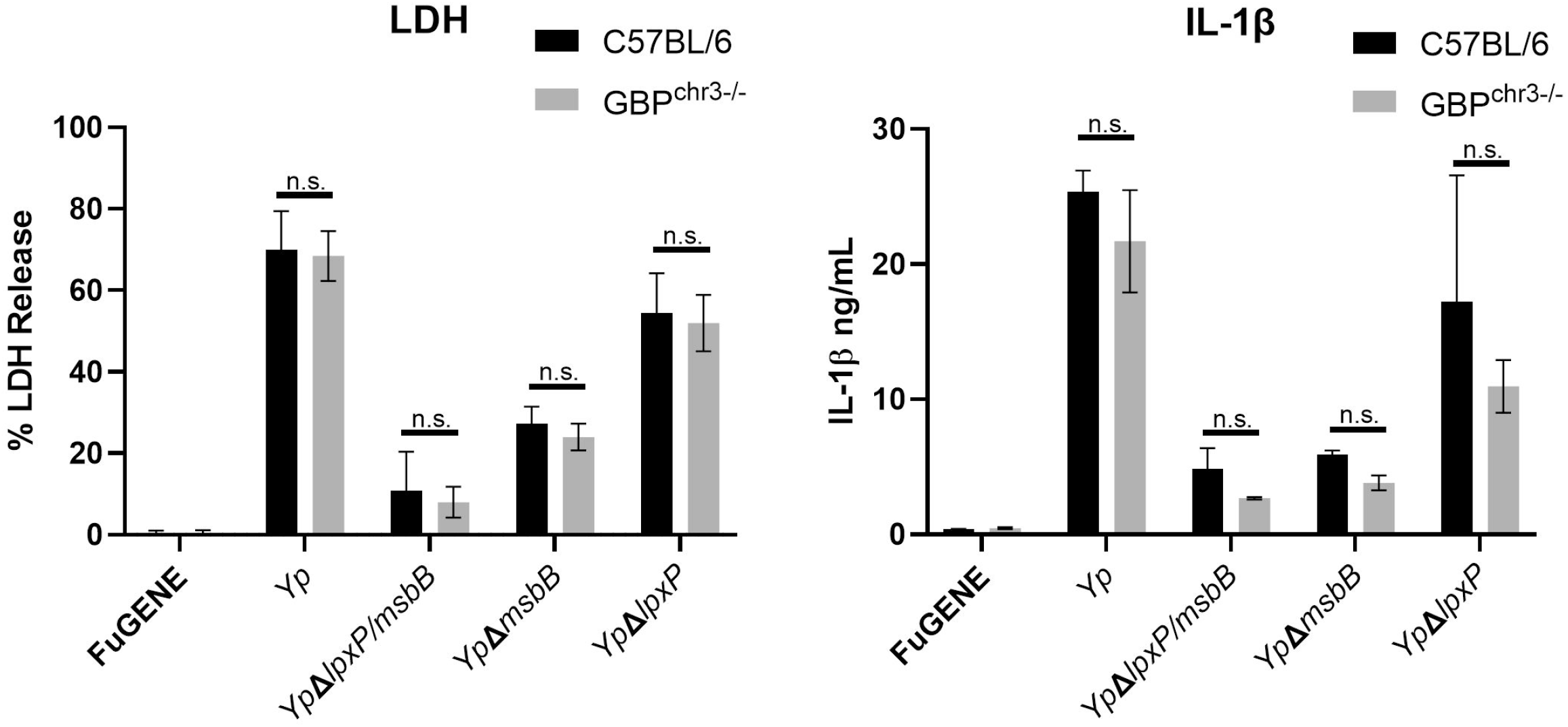
Guanylate binding proteins on Chromosome 3 are dispensable for sensing transfected intracellular LPS. Pam3CSK4 primed BMDMs from wild-type (C57BL/6) and GBP-deficient (Gbp^chr3-/-^) mice were transfected with 2µg/ml of the indicated LOS species using FuGENE. Twenty hours after transfection, cell supernatants were collected and LDH activity (A) and IL-1β concentration (B) were assayed. Data are representative of three independent experiments performed in triplicate. n.s., p>0.05, t-test using Holm-Sidak method for multiple hypothesis testing.

## Discussion

Lipid A, a major component of the outer membrane of Gram-negative bacteria, is recognized by the host innate immune system, contributing both to pathogen control and inflammatory pathology during disease. The innate immune system senses lipid A through two independent receptors, TLR4 and Casp11. While lipid A structures that are agonists or non-agonists for Casp11 and TLR4 are considered to be similar, the precise rules that determine which lipid A structures specifically activate Casp11 and how these structures are recognized are not fully understood. However, the lack of a standardized series of lipid A molecules that are uniformly isolated and prepared has limited our understanding of these processes. Here, we employed a series of defined lipid A structures derived from *Yp* to investigate the precise link between acylation state and intracellular and extracellular LPS sensing. Our results indicate that the 16:1 O-linked acyl chain of *Yp* lipid A is necessary for activation of TLR4, while the shorter 12-carbon chain at the 3’ position is mostly dispensable (**Figure 2**). Conversely, we found that the majority of Casp11 activation induced by hexa-acylated *Yp* is due to the presence of the secondary acyl chain at the 3’position of the di-glucosamine backbone (**Figures 3 and 4)**. Thus, each of these penta-acylated structures represents a minimum stimulatory lipid A structure: one for TLR4 and the other for Casp11. In the case of TLR4, differences in stimulation between the two penta-acyl structures may reflect differential binding of these molecules within the hydrophobic pocket of the MD-2 coreceptor [43]. The long 16:1-carbon chain in particular may stabilize binding, leading to enhanced receptor signaling. The structural reasons for the importance of the 3’ secondary acyl chain in Casp11 activation are currently less clear. However, this difference is not attributable to the action of GBP1, as *Gbp*^*Chr3-/-*^ BMDMs did not show a defect in their ability to respond to these lipid A structures **(Figure 5)**. Interestingly, in contrast to the reported role of GBP1 in human epithelial cells [37] and the role of GBPs in facilitating LPS sensing in the setting of bacterial infections, we did not observe a contribution of GBPs to differential Casp11 activation by *Yp* Δ*lpxP* and *Yp* Δ*msbB* lipid A. Human GBP1 has been shown to have important electrostatic interactions with the core oligosaccharide and lipid A of LPS [37]. The fact that GBP1 is necessary for intracellular LPS sensing in human but not mouse cells may explain why murine BMDMs respond to both intracellular LPS and lipid A **(Figures 3 and 4)**, whereas human macrophages respond to cytosolic LPS, but fail to detect lipid A. Indeed, when human Casp4 is introduced into mouse cells lacking Casp11, these cells are able to undergo pyroptosis in response to transfected LPS (Alexander-Floyd, Harberts, et al., companion manuscript).

Previous work on Casp11 recognition of LPS has shown that the number of acyl chains in the lipid A moiety of LPS is important for determining whether that particular structure can activate the non-canonical inflammasome. Based on these studies, it is currently thought that hepta- or hexa-acylated lipid A activate Casp11, whereas structures with fewer acyl chains do not. Unexpectedly, we identify penta-acylated lipid A structures that, despite having the same number of acyl chains, differ greatly in their ability to activate Casp11 and exhibit an inverse relationship with their ability to activate TLR4 **(Figure 6)**. Our results indicate that Casp11 recognition depends on the position and length of secondary acyl chains in the lipid A molecule. *Chlamydia* and *Rhizobium* species produce lipid A molecules that do not activate Casp11.

**Figure 6:**
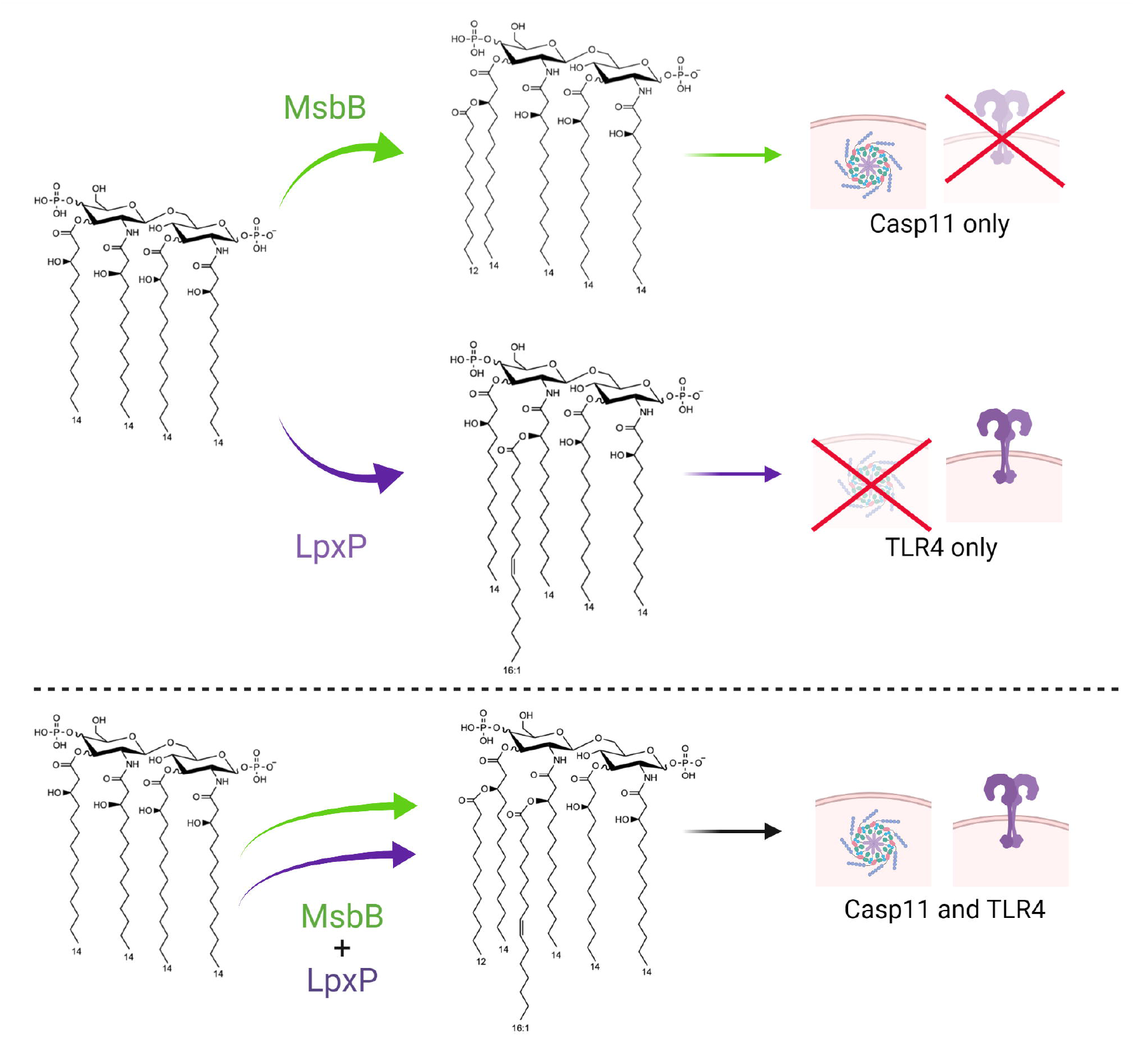
Acylation patterns of Lipid A from *Yersinia pestis* determine differential TLR4 and Caspase-11 activation [66].

Intriguingly, the structures of these lipid A molecules are similar to *Yp*Δ*msbB* in that they also lack an acyl chain at the 3’ position, implying that the presence of a secondary acyl chain at the 3’ position is a generally requirement for activation Casp11 activation [4, 23, 60-62]. A recent study that analyzed the ability of commercial preparations of *B. pertussis* LPS to activate TLR4 and Casp11 found that although *B. pertussis* LPS, which is primarily, but not exclusively, composed of penta-acylated LPS that lacks a secondary 3’ acyl chain, stimulated lower levels of IL-1α, it was still able to induce robust LDH release [53]. Our finding that mixtures of non-stimulatory and stimulatory LPS still trigger robust non-canonical inflammasome activation indicate that analysis of non-canonical inflammasome responses to lipid A require the use of lipid A preparations that contain uniform structures.

Our study reveals for the first time that distinct secondary acyl chains are critical for the activation of murine TLR4 or Casp11 by lipid A (**Figures 2-4)**. Unlike the enteric *Yersinia* species, *Y. pestis* lacks *lpxL*, which encodes a late acyltransferase that adds secondary acyl chains in other Gram-negative Enterobacteriaceae [50]. The absence of this acyltransferase, combined with the coordinate downregulation of both the *msbB* and *lpxP* genes, allows *Yp* to modify its lipid A and escape innate immune surveillance to establish a productive infection either in the arthropod or mammalian host [63]. Our findings thus suggest that the differences in LPS detection between TLR4 and Casp11 constrain the lipid A structures that can evade both components of innate immune LPS sensing. While downregulation of *lxpP* or *msbB* could allow *Yp* to evade TLR4 or Casp11, respectively, the bacterium must downregulate both to generate a tetra-acylated lipid A within a mammalian host. While *Yp* has been able to evolve such a mechanism, generation of tetra-acylated lipid A comes with a strong fitness cost. *Yp* grown at 37°C or lacking *msbB* and *lpxP* are more susceptible to cationic antimicrobial peptides [44, 49]. Indeed, other Gram-negative Enterobacteriaceae that are engineered to produce tetra-acylated LPS structures are generally attenuated for growth and highly sensitive to environmental perturbation [64]. Increased susceptibility to cationic antimicrobials and other environmental stressors for bacteria that produce under-acylated LPS molecules highlights the selective pressure that these LPS receptors place on Gram-negative pathogens.

Lipid A structures that evade both TLR4 and Casp11 have been found in non-pathogens isolated from deep sea environments[65]. However, these lipid A structures appear to avoid detection without needing to be hypo-acylated; possibly by sampling areas of lipid A chemical space that cannot be readily accessed by pathogens at the temperatures and environmental pressures found in terrestrial environments or in mammalian hosts [65]. Interestingly, mouse Casp11 and human Casp4/5 are sensitive to distinct lipid A structures, as the human non-canonical inflammasome can detect hypo-acylated lipid A species that fail to activate Casp11 [52]. Consistently, Casp4/5 do not distinguish between different species of penta-acylated lipid A in the way that Casp11 does (Alexander-Floyd, Harberts, et al., companion manuscript).

However, human TLR4 is less sensitive to hypo-acylated LPS than mouse TLR4 [30]. Altogether, our findings described here and in Alexander-Floyd *et al*. highlight the complementary recognition mechanisms displayed by both human and mouse cytosolic and cell surface LPS sensing pathways. While there are unique structures detected by all four systems, our findings collectively indicate that each single host species possesses a self-contained endotoxin sensing pathway that comprehensively surveys lipid A structures and limits the capacity of microbial pathogens to evade detection.

## Acknowledgements

We thank members of the Brodsky, Shin, and Ernst laboratories for helpful scientific discussions. We thank Drs. Masahiro Yamamoto and Dr. Boris Striepen for bone marrow from *Gbp*^*Chr3*^ mice. This work is supported in part by R01AI128530 and R01AI139102A1 (IEB), R01AI118861 and R01AI123243 (SS), HHS-NIH-NIAID-BAA2017 (RKE), T32AI095190 (EMH), T32 AR076951 (DG) and a Burroughs-Welcome Fund Investigators in the Pathogenesis of Infectious Diseases Award (IEB and SS).

## References

1. Weiss, J. and J. Barker, Diverse pro-inflammatory endotoxin recognition systems of mammalian innate immunity. F1000Res, 2018. 7.

2. Hoshino, K., et al., Cutting edge: Toll-like receptor 4 (TLR4)-deficient mice are hyporesponsive to lipopolysaccharide: evidence for TLR4 as the Lps gene product. J Immunol, 1999. 162(7): p. 3749–52.

3. Wang, S., et al., Murine caspase-11, an ICE-interacting protease, is essential for the activation of ICE. Cell, 1998. 92(4): p. 501–9.

4. Kayagaki, N., et al., Noncanonical inflammasome activation by intracellular LPS independent of TLR4. Science, 2013. 341(6151): p. 1246–9.

5. Kang, R., et al., Lipid Peroxidation Drives Gasdermin D-Mediated Pyroptosis in Lethal Polymicrobial Sepsis. Cell Host Microbe, 2018. 24(1): p. 97–108 e4.

6. Park, B.S., et al., The structural basis of lipopolysaccharide recognition by the TLR4-MD-2 complex. Nature, 2009. 458(7242): p. 1191–5.

7. Shi, J., et al., Inflammatory caspases are innate immune receptors for intracellular LPS. Nature, 2014. 514(7521): p. 187–92.

8. Xiao, X., K. Sankaranarayanan, and C. Khosla, Biosynthesis and structure-activity relationships of the lipid a family of glycolipids. Curr Opin Chem Biol, 2017. 40: p. 127–137.

9. Raetz, C.R., et al., Lipid A modification systems in gram-negative bacteria. Annu Rev Biochem, 2007. 76: p. 295–329.

10. Chilton, P.M., C.A. Embry, and T.C. Mitchell, Effects of Differences in Lipid A Structure on TLR4 Pro-Inflammatory Signaling and Inflammasome Activation. Front Immunol, 2012. 3: p. 154.

11. Park, B.S. and J.O. Lee, Recognition of lipopolysaccharide pattern by TLR4 complexes. Exp Mol Med, 2013. 45: p. e66.

12. Kawasaki, T. and T. Kawai, Toll-like receptor signaling pathways. Front Immunol, 2014. 5: p. 461.

13. Maeshima, N. and R.C. Fernandez, Recognition of lipid A variants by the TLR4-MD-2 receptor complex. Front Cell Infect Microbiol, 2013. 3: p. 3.

14. Coats, S.R., et al., MD-2 mediates the ability of tetra-acylated and penta-acylated lipopolysaccharides to antagonize Escherichia coli lipopolysaccharide at the TLR4 signaling complex. J Immunol, 2005. 175(7): p. 4490–8.

15. Steimle, A., I.B. Autenrieth, and J.S. Frick, Structure and function: Lipid A modifications in commensals and pathogens. Int J Med Microbiol, 2016. 306(5): p. 290–301.

16. Jones, B.D., et al., Study of the role of the htrB gene in Salmonella typhimurium virulence. Infect Immun, 1997. 65(11): p. 4778–83.

17. Sunshine, M.G., et al., Mutation of the htrB gene in a virulent Salmonella typhimurium strain by intergeneric transduction: strain construction and phenotypic characterization. J Bacteriol, 1997. 179(17): p. 5521–33.

18. Vaara, M. and M. Nurminen, Outer membrane permeability barrier in Escherichia coli mutants that are defective in the late acyltransferases of lipid A biosynthesis. Antimicrob Agents Chemother, 1999. 43(6): p. 1459–62.

19. Ellis, C.D., et al., The Neisseria gonorrhoeae lpxLII gene encodes for a late-functioning lauroyl acyl transferase, and a null mutation within the gene has a significant effect on the induction of acute inflammatory responses. Mol Microbiol, 2001. 42(1): p. 167–81.

20. van der Ley, P., et al., Modification of lipid A biosynthesis in Neisseria meningitidis lpxL mutants: influence on lipopolysaccharide structure, toxicity, and adjuvant activity. Infect Immun, 2001. 69(10): p. 5981–90.

21. Tong, H.H., et al., Expression of cytokine and chemokine genes by human middle ear epithelial cells induced by formalin-killed Haemophilus influenzae or its lipooligosaccharide htrB and rfaD mutants. Infect Immun, 2001. 69(6): p. 3678–84.

22. Post, D.M., et al., Intracellular survival of Neisseria gonorrhoeae in male urethral epithelial cells: importance of a hexaacyl lipid A. Infect Immun, 2002. 70(2): p. 909–20.

23. Ingram, B.O., et al., Altered lipid A structures and polymyxin hypersensitivity of Rhizobium etli mutants lacking the LpxE and LpxF phosphatases. Biochim Biophys Acta, 2010. 1801(5): p. 593–604.

24. Cullen, T.W., et al., Helicobacter pylori versus the host: remodeling of the bacterial outer membrane is required for survival in the gastric mucosa. PLoS Pathog, 2011. 7(12): p. e1002454.

25. Kong, Q., et al., Phosphate groups of lipid A are essential for Salmonella enterica serovar Typhimurium virulence and affect innate and adaptive immunity. Infect Immun, 2012. 80(9): p. 3215–24.

26. Wang, X., et al., Expression cloning and periplasmic orientation of the Francisella novicida lipid A 4’-phosphatase LpxF. J Biol Chem, 2006. 281(14): p. 9321–30.

27. Jones, J.W., et al., Comprehensive structure characterization of lipid A extracted from Yersinia pestis for determination of its phosphorylation configuration. J Am Soc Mass Spectrom, 2010. 21(5): p. 785–99.

28. Guo, L., et al., Lipid A acylation and bacterial resistance against vertebrate antimicrobial peptides. Cell, 1998. 95(2): p. 189–98.

29. Coats, S.R., et al., Human Toll-like receptor 4 responses to P. gingivalis are regulated by lipid A 1-and 4’-phosphatase activities. Cell Microbiol, 2009. 11(11): p. 1587–99.

30. Vaure, C. and Y. Liu, A comparative review of toll-like receptor 4 expression and functionality in different animal species. Front Immunol, 2014. 5: p. 316.

31. Hajjar, A.M., et al., Human Toll-like receptor 4 recognizes host-specific LPS modifications. Nat Immunol, 2002. 3(4): p. 354–9.

32. Kawahara, K., et al., Modification of the structure and activity of lipid A in Yersinia pestis lipopolysaccharide by growth temperature. Infect Immun, 2002. 70(8): p. 4092–8.

33. Aachoui, Y., et al., Caspase-11 protects against bacteria that escape the vacuole. Science, 2013. 339(6122): p. 975–8.

34. Hagar, J.A., et al., Cytoplasmic LPS activates caspase-11: implications in TLR4-independent endotoxic shock. Science, 2013. 341(6151): p. 1250–3.

35. Kayagaki, N., et al., Non-canonical inflammasome activation targets caspase-11. Nature, 2011. 479(7371): p. 117–21.

36. Meunier, E., et al., Caspase-11 activation requires lysis of pathogen-containing vacuoles by IFN-induced GTPases. Nature, 2014. 509(7500): p. 366–70.

37. Santos, J.C., et al., Human GBP1 binds LPS to initiate assembly of a caspase-4 activating platform on cytosolic bacteria. Nat Commun, 2020. 11(1): p. 3276.

38. Pilla, D.M., et al., Guanylate binding proteins promote caspase-11-dependent pyroptosis in response to cytoplasmic LPS. Proc Natl Acad Sci U S A, 2014. 111(16): p. 6046–51.

39. Wandel, M.P., et al., Guanylate-binding proteins convert cytosolic bacteria into caspase-4 signaling platforms. Nat Immunol, 2020. 21(8): p. 880–891.

40. Lee, B.L., et al., Caspase-11 auto-proteolysis is crucial for noncanonical inflammasome activation. J Exp Med, 2018. 215(9): p. 2279–2288.

41. Kayagaki, N., et al., Caspase-11 cleaves gasdermin D for non-canonical inflammasome signalling. Nature, 2015. 526(7575): p. 666–71.

42. Shi, J., et al., Cleavage of GSDMD by inflammatory caspases determines pyroptotic cell death. Nature, 2015. 526(7575): p. 660–5.

43. Kim, H.M., et al., Crystal structure of the TLR4-MD-2 complex with bound endotoxin antagonist Eritoran. Cell, 2007. 130(5): p. 906–17.

44. Rebeil, R., et al., Characterization of late acyltransferase genes of Yersinia pestis and their role in temperature-dependent lipid A variation. J Bacteriol, 2006. 188(4): p. 1381–8.

45. Gregg, K.A., et al., Rationally Designed TLR4 Ligands for Vaccine Adjuvant Discovery. mBio, 2017. 8(3).

46. Hirschfeld, M., et al., Cutting edge: repurification of lipopolysaccharide eliminates signaling through both human and murine toll-like receptor 2. J Immunol, 2000. 165(2): p. 618–22.

47. Scott, A.J., et al., Lipid A structural modifications in extreme conditions and identification of unique modifying enzymes to define the Toll-like receptor 4 structure-activity relationship. Biochim Biophys Acta Mol Cell Biol Lipids, 2017. 1862(11): p. 1439–1450.

48. Vadyvaloo, V., et al., Transit through the flea vector induces a pretransmission innate immunity resistance phenotype in Yersinia pestis. PLoS Pathog, 2010. 6(2): p. e1000783.

49. Rebeil, R., et al., Variation in lipid A structure in the pathogenic yersiniae. Mol Microbiol, 2004. 52(5): p. 1363–73.

50. Montminy, S.W., et al., Virulence factors of Yersinia pestis are overcome by a strong lipopolysaccharide response. Nat Immunol, 2006. 7(10): p. 1066–73.

51. Hajjar, A.M., et al., Humanized TLR4/MD-2 mice reveal LPS recognition differentially impacts susceptibility to Yersinia pestis and Salmonella enterica. PLoS Pathog, 2012. 8(10): p. e1002963.

52. Lagrange, B., et al., Human caspase-4 detects tetra-acylated LPS and cytosolic Francisella and functions differently from murine caspase-11. Nat Commun, 2018. 9(1): p. 242.

53. Ernst, O., et al., Species-Specific Endotoxin Stimulus Determines Toll-Like Receptor 4-and Caspase 11-Mediated Pathway Activation Characteristics. mSystems, 2021. 6(4): p. e0030621.

54. Brubaker, S.W., et al., A Rapid Caspase-11 Response Induced by IFN-gamma is Independent of Guanylate Binding Proteins. iScience, 2020. 23(10): p. 101612.

55. Feeley, E.M., et al., Galectin-3 directs antimicrobial guanylate binding proteins to vacuoles furnished with bacterial secretion systems. Proc Natl Acad Sci U S A, 2017. 114(9): p. E1698–E1706.

56. Zwack, E.E., et al., Guanylate Binding Proteins Regulate Inflammasome Activation in Response to Hyperinjected Yersinia Translocon Components. Infect Immun, 2017. 85(10).

57. Fisch, D., et al., Human GBP1 is a microbe-specific gatekeeper of macrophage apoptosis and pyroptosis. EMBO J, 2019. 38(13): p. e100926.

58. Man, S.M., et al., Interferon-inducible guanylate-binding proteins at the interface of cell-autonomous immunity and inflammasome activation. J Leukoc Biol, 2017. 101(1): p. 143–150.

59. Yamamoto, M., et al., A cluster of interferon-γ-inducible p65 GTPases plays a critical role in host defense against Toxoplasma gondii. Immunity, 2012. 37(2): p. 302–13.

60. Muszynski, A., et al., Structures of the lipopolysaccharides from Rhizobium leguminosarum RBL5523 and its UDP-glucose dehydrogenase mutant (exo5). Glycobiology, 2011. 21(1): p. 55–68.

61. Yang, C., et al., Chlamydia trachomatis Lipopolysaccharide Evades the Canonical and Noncanonical Inflammatory Pathways To Subvert Innate Immunity. mBio, 2019. 10(2).

62. Alexander, J., et al., Manuscript co-submitted. 2021.

63. Demeure, C.E., et al., Yersinia pestis and plague: an updated view on evolution, virulence determinants, immune subversion, vaccination, and diagnostics. Genes Immun, 2019. 20(5): p. 357–370.

64. Vorachek-Warren, M.K., et al., A triple mutant of Escherichia coli lacking secondary acyl chains on lipid A. J Biol Chem, 2002. 277(16): p. 14194–205.

65. Gauthier, A.E., et al., Deep-sea microbes as tools to refine the rules of innate immune pattern recognition. Sci Immunol, 2021. 6(57).

66. “Created with BioRender.com”.

