## Supplemental Materials for "Position-specific secondary acylation determines detection of lipid A by murine TLR4 and caspase-11"

**Supplemental Figure 1: Delivery of LPS using *L. monocytogenes* highlights the role of acyl chain position in Casp11 activation.** Wild-type or Casp11$^{-/-}$ BMDMs were primed with poly(I:C) and then infected with *L. monocytogenes* in the presence of LPS/LOS. Four hours after infection, cell death was assayed by LDH release assay. Data are presented as mean percentage cell death across technical replicates ± SD and are representative of three independent experiments. *, significant difference between C57BL/6 and Casp11$^{-/-}$, p<0.05, t-test using Holm-Sidak method for multiple hypothesis testing.

**Supplemental Figure 2: Penta-acyl LPS molecules differ in their ability to inhibit Casp11.** BMDMs were primed with Pam3CSK4 and then transfected with 1µg/ml LPS from *Salmonella* Minnesota (LPS-*Sm*) alone or with increasing doses of one of the indicated hypo-acylated LPS molecules. (LPS-*Rs* indicates LPS from *Rhodobacter sphaeroides*) Twenty hours after transfection, cell supernatants were assayed for lactate dehydrogenase activity. Lactate dehydrogenase activity was determined relative to cells lysed using Triton X-100. Data are presented as averages (+/- SD) of three independent experiments performed in triplicate. *, significant difference from LPS-*Sm* alone, p<0.05, One-way ANOVA with Dunnet's multiple comparisons test.

**Supplemental Figure 1.**

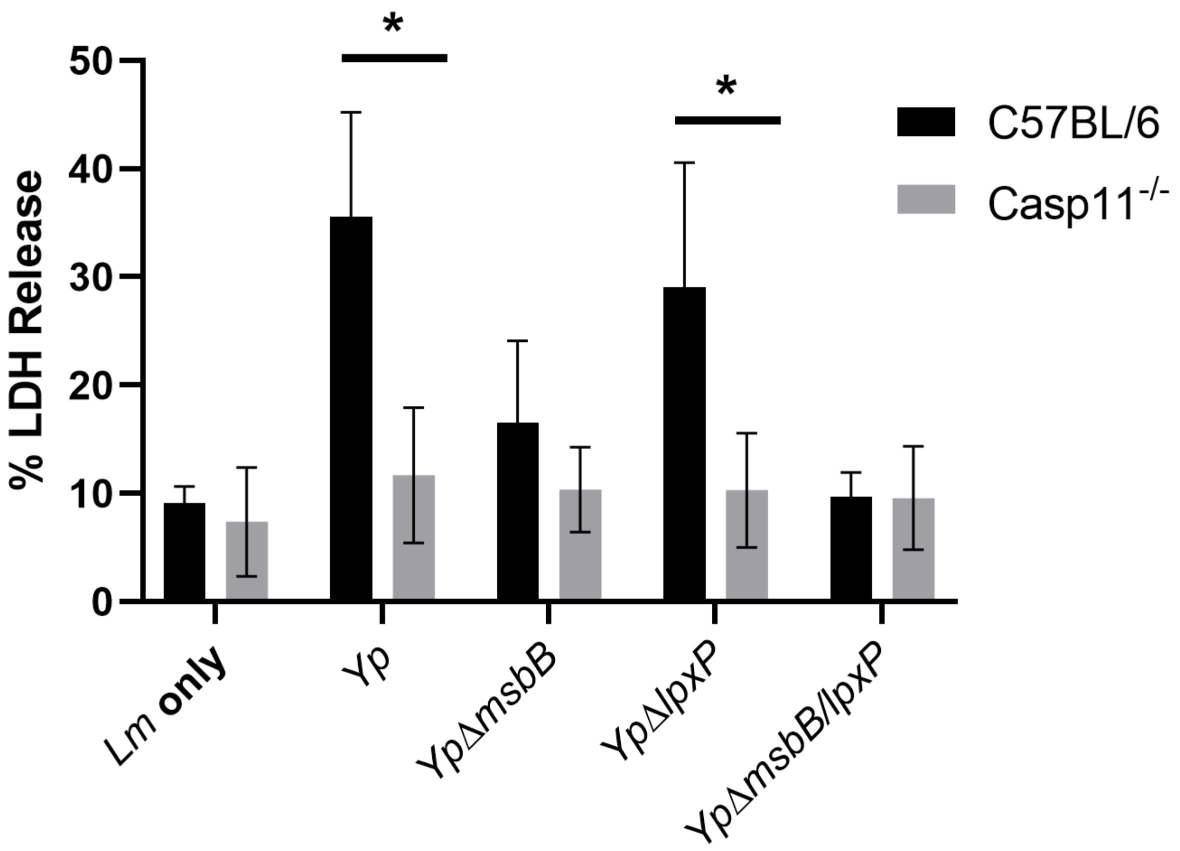

**Supplemental Figure 2.**

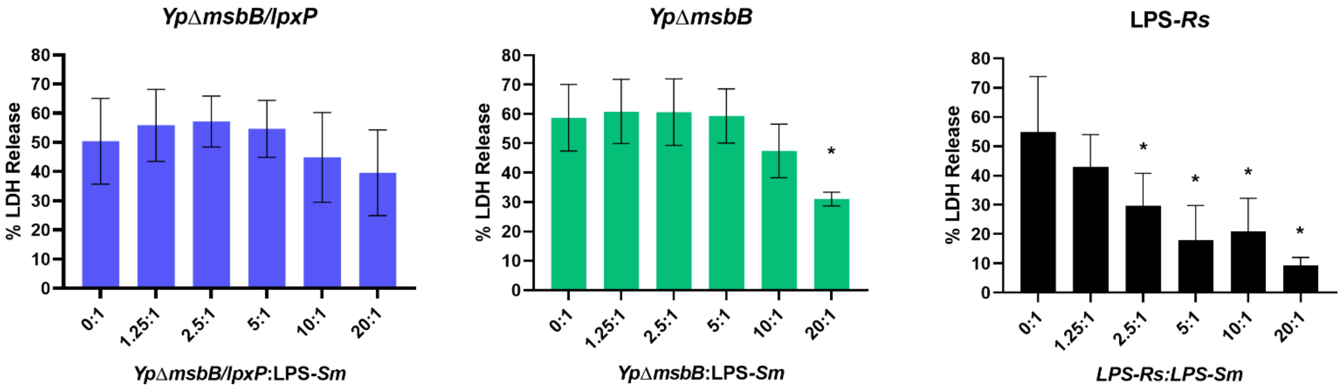
